# Kv2.1-Kv6.4 subunits deficiency impairs inhibitory signaling and visual circuit dynamics in zebrafish

**DOI:** 10.1101/2025.10.30.685505

**Authors:** Ruchi P Jain, Rosa R Amini, Lukasz Majewski, Vladimir Korzh

## Abstract

Voltage-gated potassium channels (Kv) play a crucial role in maintaining the cell’s resting potential. Mutations in the Kv2.1 voltage-gated potassium channel are associated with developmental epileptic encephalopathy. Previous analysis of the loss-of-function zebrafish mutants *kcnb1^sq301^* and *kcng4b^waw304^,* which affect genes that encode Kv2.1 subunits (the α subunit Kcnb1 and the modulatory subunit Kcng4b), has shown that they play an antagonistic role in the development of hollow organs, such as the brain and ear. In this study, we investigated the behavioral effects of these mutations. Under normal light conditions, both mutants exhibited reduced activity at 5 days post-fertilization. However, exposure to a low concentration of 5 mM of the chemoconvulsant pentylenetetrazole increased their locomotor activity and induced seizures. Quantitative RT-PCR (qRT-PCR) analysis of the mutants revealed an increase in the transcript levels of *c-fos* and *gad2,* and a decrease in a transcript level of *gabra1.* This suggests that mutations cause a disruption to inhibitory neurotransmission. Local field potential recordings from the optic tectum of the mutants under baseline conditions showed an increase in spontaneous electrical activity. The *kcnb1* mutant is photosensitive and experiences freezing episodes under high-intensity light. Together, these findings suggest that defects in Kv2.1 subunits impact both locomotor behavior and light-evoked responses.

## INTRODUCTION

Voltage-gated potassium channels facilitate the re-polarization of the action potential (1–3). Consequently, genetic mutations in these channels are a frequent cause of severe neurological disorders (4–7). Among these, mutation in the electrically active subunit Kv2.1 gene, encoded by *KCNB1*, has been linked to a spectrum of conditions, including developmental epileptic encephalopathy (DEE) (8–11). More than 50 *de novo* pathogenic variants in the *KCNB1* gene have been identified in DEE patients. The patients suffer from unprovoked seizures, intellectual disability, autism spectrum disorder (ASD), and often severe movement disorders (12–18). Kv2.1 is predominantly localized to the somatodendritic region of the central neurons, where it generates a major component of the delayed rectifier K^+^ current that is critical for shaping the action potential duration and regulating overall excitability (19, 20). Therefore, these mutations alter the channel’s gating properties or its membrane localization due to the gain-of-function (GOF), loss-of-function (LOF), and dominant negative effects, i.e. the changes that disturb the intricate balance of excitation and inhibition within the neural circuits (21–23). However, despite the number of cases of *KCNB1*-related DEE increases, the mechanism by which Kv2.1 dysfunction influences circuit function and sensory-driven behavior during early brain development remains poorly understood.

Kv2.1 often forms heterotetrameric complexes with electrically silent/modulatory subunits KvS, such as Kv5, Kv6, Kv8, and Kv9, which do not form functional channels independently, but instead co-assemble with Kv2.1 to modulate its biophysical and functional properties (24–26). One such silent subunit is Kv6.4 (encoded by *KCNG4*), a key partner that imposes distinct regulatory effects. Kv6.4’s co-assembly in the mammalian central nervous system drastically hyperpolarizes the inactivation kinetics of Kv2.1 by approximately -40 mV (27,28). In developing zebrafish, *kcnb1* and *kcng4b* show overlapping expression patterns in the brain, suggesting that the Kv2.1 function is conserved in evolution. These genes play a role in the development of hollow organs like the brain ventricles and inner ear, and an assembly of Reissner Fiber. This highlights the developmental aspect of Kv2.1/Kv6.4 interactions extending beyond their canonical role characterized by electrophysiology studies (29–32). Conversely, in the mammalian photoreceptors, Kv2.1 forms heterotetrameric channels with Kv8.2, encoded by *KCNV2*. This assembly is important as the presence of Kv8.2 shifts the voltage-dependence of Kv2.1, generating a continuous ‘window current’ that is vital for regulating the photoreceptor’s dark current and subsequent signal transmission (33,34).

Despite this clear link between *KCNB1* and severe DEE, the precise mechanisms by which Kv2.1 loss leads to complex phenotypes remain poorly understood. Specifically, there are two major gaps: First, while Kv2.1’s role in central neuronal hyperexcitability is accepted, the contribution of its modulatory subunit, Kv6.4/KCNG4, to the shared DEE pathophysiology has not been rigorously compared in a unified model. Second, a minority of *KCNB1* patients exhibit photosensitive symptoms (such as myoclonic eyelid seizures) (35), yet the specific mechanism by which Kv2.1 regulates light-evoked behavior and whether this defect originates in the brain or retina remains unknown.

To address these gaps, we used the direct, comparative assessment of the neural functions and behavior of the zebrafish LOF mutants *kcnb1^-/-^* with defect in the electrically active α-subunit (Kcnb1), and *kcng4b^-/-^* with defect in the silent modulatory subunit (Kcng4b).

In this paper, we report two principal findings that define the neurological and sensory consequences of subunit deficiency. Firstly, we demonstrate that both *kcnb1^-/-^* and *kcng4b^-/-^* mutants share a pathophysiology of central neuronal hyperexcitability, characterized by increased spontaneous brain activity, elevated *c-fos* expression, and increased susceptibility to pentylenetetrazole-induced seizures. Furthermore, we reveal a compensatory mechanism involving the inhibitory neurotransmitter system (*gad2* and *gabra1*) that underlies this hyperexcitable state. Secondly, we identify a unique defect in *kcnb1^-/-^* mutants - a profound and acute increase in sensitivity to light and freezing behavior. We show that this behavioral phenotype is not caused by structural defects in the retina. Most likely, it is the result of a ‘two-hit’ mechanism where a subtle signaling defect originating in the photoreceptors is amplified by the central neural hyperexcitability. Collectively, our work establishes shared and distinct roles for the two Kv2.1’ subunits, highlighting their essential contribution to both inhibitory circuit homeostasis and sensory processing.

## MATERIAL AND METHODS

### Animals

We used three groups of zebrafish (Danio rerio), including a wild type (AB) control (WT), the *kcnb1* null mutant *kcnb1^sq301^* (29), and the *kcng4b* mutant *kcng4b^waw304^* (36). Only the homozygous mutants (*kcnb1^-/-^* and *kcng4b^-/-^*) were used. However, for the readability throughout the manuscript, we use the gene shorthand *kcnb1* and *kcng4b* to refer to these mutants, respectively. The fish were maintained following established protocols (37) at the Zebrafish Core Facility of the International Institute of Molecular and Cell Biology, Warsaw (License number PL14656251). Procedures involving embryonic and larval zebrafish were conducted per the guidelines set by the Polish Laboratory Animal Science Association and the Council Directive (63/2010/EEC) of the European Communities.

### Behavioral experiments

All behavioral experiments were conducted on 5 days post fertilization (dpf) larvae using the Zebrabox system (Viewpoint, Life Sciences, France) and Ethovision XT 12 software (Noldus) for tracking and analysis. All experiments were repeated at least three times. The number of larvae used for each experiment were specified in the figure legends. Locomotor activity was quantified by calculating the total distance travelled and mean velocity.

### **a.** Pentylenetetrazole (PTZ) induced seizures

4 dpf larvae were transferred to a 96-well plate with 100 μl of E3 medium acclimatized overnight at 28.5°C. The plate was then placed in the Zebrabox for a 30 min dark acclimatization period. Baseline behavior was recorded with 50% bottom light for 15 min. Subsequently, 100 μl of fresh PTZ solution (#P6500, Sigma) at various concentrations (10, 20, and 30 mM) was added to the wells, resulting in final concentrations 5, 10, and 15 mM. Control wells received 100 μl of E3 medium and recorded for 30 min. Hyperactivity was determined by comparing total distance and mean velocity before and after PTZ treatment.

### **b.** Dual light pulse-induced seizures

4 dpf larvae were acclimatized overnight at 28.5°C in a 96-well plate with 200 μl of E3 medium. On the next day, larvae were dark acclimatized for a 30 min period in the Zebrabox, followed by a 15 min baseline recording. To induce seizures, two light pulses (0.5 s each-100% bottom light intensity) separated by 1 s of darkness were administered every 2 minutes for a total of 10 min (38). To assess photosensitivity, the total distance and mean velocity were calculated.

### **c.** Light-dark preference

To assess light-dark preference, we used a 10 cm Petri dish. Half of the plate was designated as dark zone by covering its side with black tape and its top half with a photographic filter (Cokin P154 ND8, Rungis, France) to block the overhead light. The other half was designated as light zone. A single 5-dpf larva was placed in the dish containing 20 ml of E3 medium and transferred to the Zebrabox system with 70% top light intensity. The experiment started after 2 min setup period and locomotor activity was recorded for 15 min. The preference was quantified by calculating the total time spent in each zone, the total distance traveled, mean velocity and frequency (39). Heat maps were obtained using the following Ethovision settings: merging method - mean, smoothing - 5, and map colors per heat map.

### **d.** Open field test

Individual larvae were transferred to a 24-well plate and placed in the Zebrabox system. For dark conditions, the entire experiment was performed in complete darkness (light intensity = 0%), whereas for light conditions, the experiment was performed with 70% top light intensity. The locomotor activity was recorded for 15 mins, following a 2 min acclimatization. To determine the locomotor behavior under different light conditions, the total distance and mean velocity were calculated. Heat maps were generated using the same settings as described in the light-dark preference test.

### **e.** Light intensity test

This protocol was developed to assess the larvae’s tolerance to varying light intensity. Larvae were individually transferred to a 24-well plate and placed inside the Zebrabox system in 100% darkness for 2 min before the experiment. Locomotor activity was recorded for 12 min while top light intensity was gradually increased by 20% every 2 minutes, starting from 0 to 100% and returning to 0% at the 12^th^ minute. Light intensity tolerance was determined by calculating the total distance, mean velocity, and freezing activity.

### **f.** Behavioral data analysis and statistics

For all behavioral experiments, the raw data were processed using Ethovision XT 12 and exported as Microsoft Excel files. Statistical analysis was performed using GraphPad Prism (Version 9.5.1). Normality of data distribution was assessed using the Shapiro-Wilk test. Depending on the outcome, comparisons were done using appropriate parametric (unpaired Student’s *t*-test, one-way ANOVA, or two-way ANOVA followed by a post-hoc test) or non-parametric (Mann-Whitney *U* test or Kruskal-Wallis test followed by a post-hoc test). A p-value of ≤0.05 was considered statistically significant.

### Quantitative Polymerase chain reaction (qPCR) analysis

qRT-PCR was performed to quantify the expression of genes related to epilepsy and photoreceptors. RNA was isolated from 50 whole 5 dpf larvae using TRIzol - chloroform (#T9424, Sigma) method according to the manufacturer’s protocol. RNA concentration was quantified, and 1 μg of RNA was reverse-transcribed into cDNA using the iScript cDNA synthesis kit (Biorad). qRT-PCR and subsequent data analysis were performed as previously described (31). A list of genes with their primer sequences are provided in the supplementary table 1.

### Immunohistochemistry

For staining sections, 5 dpf larvae were fixed in 4% PFA (#P6148, Sigma)/PBS overnight. After PBS washes, they were fixed in 1.5% bactoagar (#214010, BD Biosciences) mixed with 5% sucrose (#772090110, Avantor)/PBS followed by transfer to a 30% sucrose solution for overnight incubation at 4°C until they sank. The bactoagar-embedded larvae were mounted in OCT (#361603E, VWR) and sectioned (14-18 μm) using a Leica cryostat. Sections were allowed to dry overnight and processed for staining as per (40). Primary antibodies used were rabbit polyclonal anti-gnat2 (zebrafish) (1:500) (#PM075, MBL) for cone photoreceptors, and mouse anti-glutamine synthetase (#610517, BD Biosciences) for Müller glia. Following PBSTx (1x PBS+ 0.1 % Triton-X 100) washes, embryos were incubated in secondary antibody Alexa Fluor anti-mouse 594 or anti-rabbit 594 (1:1000) for 2 hr. Hoechst (#H3570, Invitrogen) was used for nuclear staining.

### Retinotectal projection tracing via Dil labelling

Phenylthiourea (PTU, #L06690, ThermoFisher) treated 5 dpf larvae were fixed overnight in 4% PFA/PBS at 4°C, followed by three 1-hour PBS washes. Fixed larvae were embedded in 1.5% low-melting agarose (LMA). Anterograde labelling of retinal ganglion cell axons was performed by microinjecting 5mg/ml DiI (1,1’-Dioctadecyl-3,3,3’,3’-Tetramethylindocarbocyanine Perchlorate, #D282, Molecular Probes), diluted in dimethyl sulfoxide, into the anterior retina. Embryos were then left overnight at room temperature. Injected embryos were visualized under Zeiss Lightsheet Z1 microscope using a 40x lens.

### Local field potential recording

Five-day-old larva was initially paralyzed using 300 μg of pancuronium bromide (#P1918, Sigma) and then embedded in 1.5% low melting point agarose (LMA, #50080, Lonza) made in embryo medium containing pancuronium bromide, on the sample holder. The sample holder was then placed on the stage of a Zeiss Axio Examiner Z1 confocal microscope and filled with freshly prepared ACSF (artificial cerebrospinal fluid). A glass microelectrode (2-7 MΩ) was backloaded with ACSF and tactically inserted into the optic tectum. Spontaneous electrical activity was recorded for 15 min using a MultiClamp 700B amplifier (Axon Instruments, United States). The resulting voltage records were digitized at a sampling rate of 20 kHz using Digidata 1550B (Axon Instruments, United States), and band-pass filtered at 0.5 Hz (high-pass) and 0.3 kHz (low-pass).

### Electrophysiology data analysis and statistics

The recordings were processed and analyzed using custom scripts written in Python (Version 3.13.5). Analysis included bandpass filtering, adaptive thresholding, and quantification of spontaneous transient events (STE) by amplitude and duration. The script is freely available upon request from the corresponding author.

The resulting summary data were analyzed for statistical significance using GraphPad Prism (Version 9.5.1). Following the Shapiro-Wilk test for assessing data normality, statistics were calculated using one-way ANOVA followed by Dunnett’s multiple comparison test. A p-value of ≤0.05 was considered statistically significant.

## RESULTS

### Kv2.1-Kv6.4 subunit deficiency does not cause spontaneous seizure but alters baseline locomotion

To assess for unprovoked seizure activity, which is a characteristic of KCNB1-related DEE (13), we recorded the baseline locomotor behavior of larvae under standard light conditions (50% bottom light). Deficiency of Kv2.1 and Kv6.4 subunits did not result in spontaneous seizure activity. However, the mutants exhibited reduced baseline locomotor activity compared to the wild-type embryos (Fig. 1a and b).

**Fig. 1:**
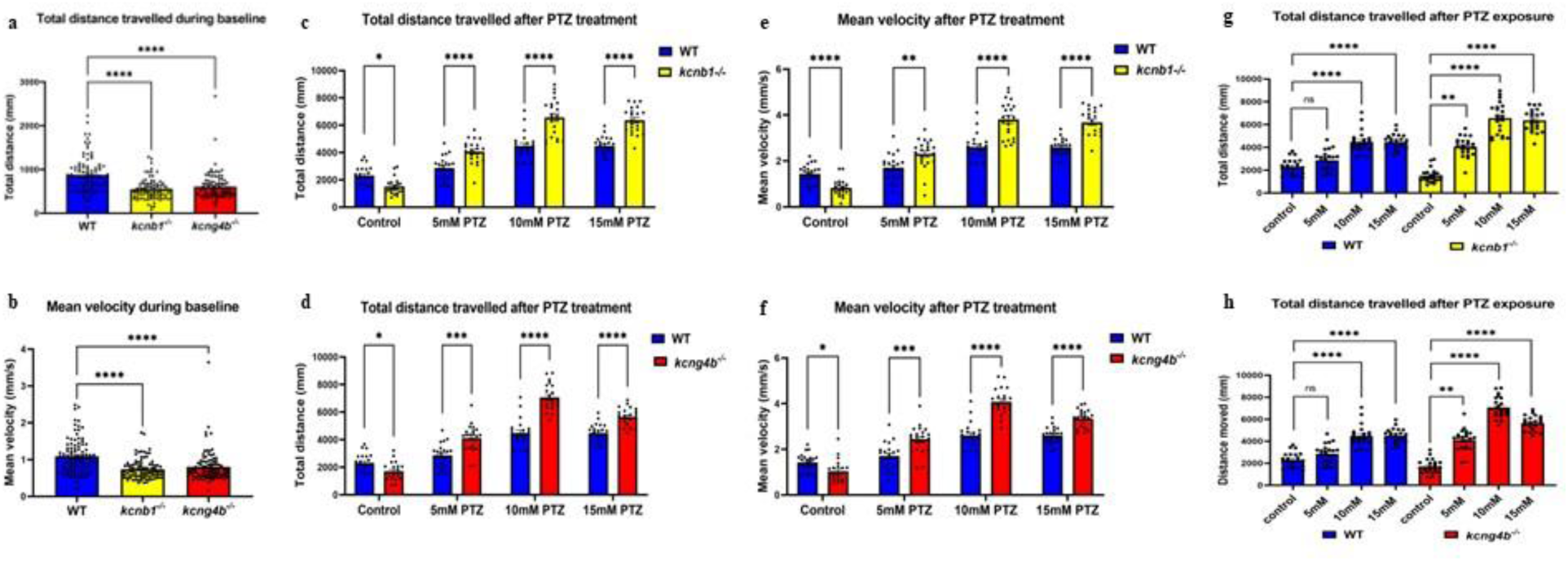
Kv2.1 and Kv6.4 subunit mutants are more susceptible to PTZ-induced generalized seizures. All experiments were performed at 5 dpf with 50% bottom light intensity in Zebrabox. **a-b) Baseline motor activity:** Baseline recording for 15 min shows decreased total distance (a) and mean velocity (b) in the *kcnb1^-/-^* and *kcng4b^-/-^* larvae compared to WT. Statistics calculated using non-parametric Kruskal-Wallis test followed by Dunn’s multiple comparison test. (N=3; n WT=93, n *kcnb1^-/-^*=93, n *kcng4b^-/-^*=89); **c-f) Concentration-dependent hyperactivity:** The total distance traveled and mean velocity are significantly increased in the *kcnb1* mutant larvae (c and e) and *kcng4b* mutant larvae (d and f) after treatment with different concentrations (5, 10 and 15 mM) of pentylenetetrazole (PTZ) for 30 min. Statistics were calculated using two-way ANOVA followed by Ŝídák’s multiple comparison test. (N=3; n=22 for each concentration); **g-h) Seizure susceptibility at low concentration:** A significant increase in the total distance travelled by both *kcnb1^-/-^* and *kcng4b^-/-^* larvae was observed at the lowest used PTZ concentration (5 mM). Statistics were calculated using non-parametric Kruskal-Wallis test followed by Dunn’s multiple comparison test. (N=3; n=22 for each concentration). All data are represented as mean ± SEM with each dot representing individual larvae. A p-value of ≤0.05 was considered statistically significant.

### Increased susceptibility to PTZ-induced seizures

We then evaluated the seizure susceptibility of the mutant larvae using PTZ, a non-competitive GABA antagonist that binds to the postsynaptic GABA-A receptor, resulting in reduced chloride ion conductance, thereby decreasing inhibitory response (41). Zebrafish have been a well-established model to study PTZ-induced seizure for the past two decades (42). We performed a dose-dependent behavioral study using PTZ concentrations of 5, 10, and 15 mM. Both WT and the mutant (*kcnb1* and *kcng4b*) larvae exhibited a dose-dependent increase in locomotor activity, as measured by total distance travelled (Fig. 1c and d) and mean velocity (Fig. 1e and f) across the different concentrations. Critically, the magnitude of hyperactivity was significantly greater in both the subunit mutants compared to the WT larvae. The seizure threshold was lower in the mutants, as 5 mM PTZ showed a significant increase in the locomotor activity of the mutants but not in the WT (Fig. 1g and h). Overall, these findings suggest that Kv2.1 and Kv6.4 subunit mutants are more susceptible to PTZ-induced generalized seizures.

### Increased spontaneous brain activity

To understand the effect of Kv2.1-Kv6.4 deficiency on electrical activity, we recorded local field potential from the optic tectum of 5 dpf larval zebrafish (Fig. 2a). Analysis of the recordings showed that spontaneous brain activity was significantly increased in both the mutants compared to WT, as evidenced by increase in total event count (Fig. 2c) and event frequency/min (Fig. 2d). The median half-width was also significantly higher in *kcnb1* mutant (Fig. 2e). Collectively, these data demonstrate that the loss of subunits leads to a significant increase in spontaneous electrical activity within the optic tectum, suggesting an underlying state of neural hyperexcitability.

**Fig. 2:**
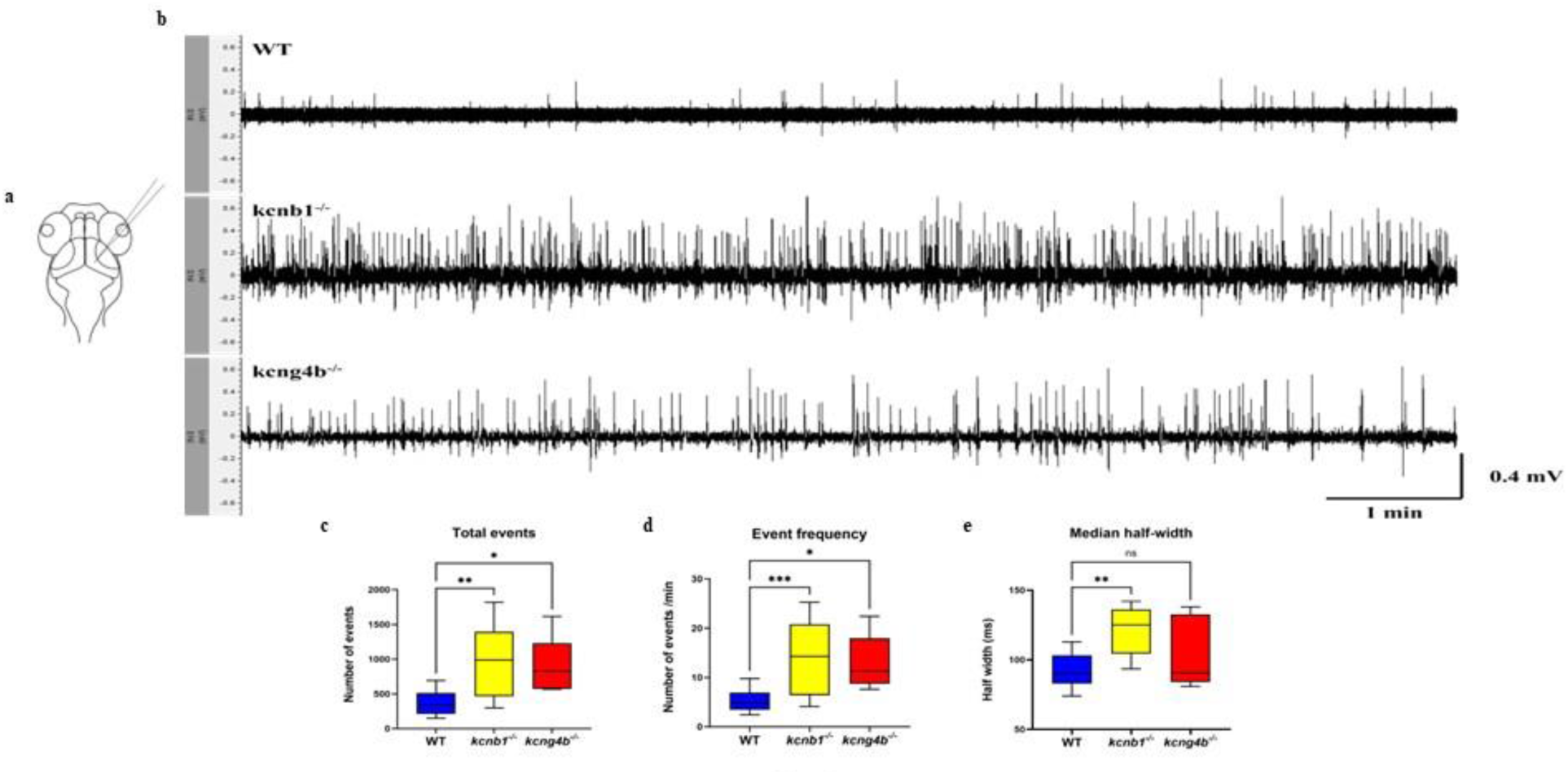
Increased spontaneous brain activity in *kcnb1^-/-^* and *kcng4b^-/-^* larva. Local field potential (LFP) was recorded from the optic tectum of 5 dpf larvae. a) Schematic representation of LFP recording site in the larval zebrafish brain. b) Representative LFP traces from a WT, *kcnb1^-/-^* and *kcng4b^-/-^* larva, showing spontaneous brain activity over a 15-minute period. **c-e) Quantitative analysis of spontaneous events:** The number of total events (c), frequency of events (d), and median half width (e) were significantly increased in the *kcnb1* mutant. The *kcng4b* mutants also showed significant increase in total events and event frequency compared to wildtype. The data are represented as box plots. The central line of box plot represents median while the whiskers extend to the minimum and maximum value of distribution. Statistics were calculated using one-way ANOVA followed by Dunnett’s multiple comparison test. Number of larvae: WT=12; *kcnb1^-/-^*=12; *kcng4b^-/-^*=5. A p-value of ≤0.05 was considered statistically significant.

### Kv2.1 deficiency altered locomotor activity triggered by dual light pulse

To determine if the Kv2.1 subunit mutants exhibit photosensitive seizures, a characteristic feature of Jeavons syndrome (43), we subjected the larvae to a dual light pulse protocol (100% - bottom light, two 0.5 s flashes separated by 1 s of darkness). Under baseline dark conditions, the overall locomotor activity of the mutant larvae was comparable to that of wild-type (Fig. 3a and b). However, during the dual light pulse test, the locomotor activity of *kcnb1^-/-^* larvae was significantly decreased, evidenced by reduced total distance and mean velocity (Fig. 3c and d). In contrast, the *kcng4b* mutant larvae behaved similarly to wild-type controls, except for a short increase in activity observed after the initial dual pulse (Fig. 3e and f). Although neither mutant displayed overt photosensitive seizures, the significant reduction in locomotor activity during the dual light pulse test indicates an altered photomotor response in the *kcnb1* mutant.

**Fig. 3:**
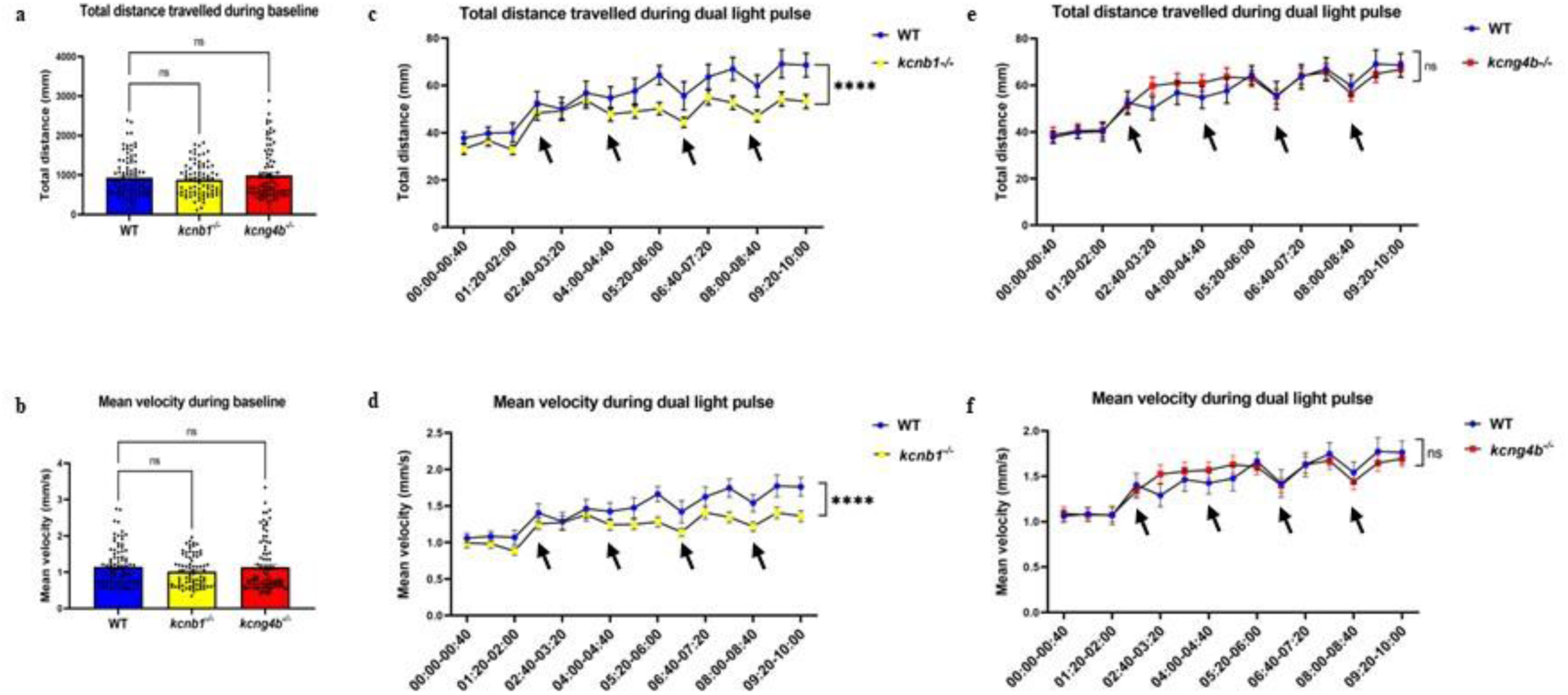
The kcnb1^-/-^ larvae are sensitive to light. The dual light pulse test was performed at 5 dpf with 100% bottom light. Red arrows mark the time when dual light pulse was flashed. **a-b) Baseline locomotor activity:** Baseline recording for 15 min in the dark showed no changes in the locomotor behaviour of *kcnb1^-/-^* and *kcng4b^-/-^* larvae as assessed by total distance (a) and mean velocity (b). Statistics were calculated using non-parametric Kruskal-Wallis followed by Dunn’s multiple comparison test. (N=3; n WT=92, n *kcnb1^-/-^* =85, n *kcng4b^-/-^*=88); **c-f) Dual light pulse response:** The overall distance traveled (c) and the mean velocity (d) are significantly decreased in the *kcnb1* mutant larvae during dual light pulse experiment suggesting light sensitivity; Conversely, no significant changes were observed in the locomotor activity of *kcng4b* mutants (e-f) under same conditions. Statistics were calculated via non-parametric Kruskal-Wallis followed by Dunn’s multiple comparison test. (N=3; n WT=68, n *kcnb1^-/-^*=85, n *kcng4b^-/-^*=88). All data are represented as mean ± SEM with each dot representing individual larvae. A p-value of ≤0.05 was considered statistically significant.

### *Kcnb1* LOF mutants exhibit photosensitivity to high-intensity light

To further evaluate the light-sensitive behavior of *kcnb1* mutants, a light/dark preference test was performed in a Petri dish with 70% top light; half of the dish was covered using a photographic film to make the dark zone (Fig. 4a). Both the wild-type and *kcnb1* mutant larvae spent more time and covered larger distances in the light zone than in the dark, while the *kcng4b* mutant showed no significant preference for either zone (Fig. 4b and c). However, the mean velocity difference between the wildtype and mutant larva remained unaltered (Supplementary Fig. 1a), whereas the zone transition frequency increased in the light zone for WT and *kcnb1* mutant, but not in the *kcng4b* mutant (Supplementary Fig. 1b). On the other hand, the heatmaps of 14/32 of the *kcnb1* mutant larvae exhibited freezing bouts (Fig. 4d) after being exposed to high-intensity light, while only 2/32 wild-type and 5/30 *kcng4b* mutant larvae froze in the same conditions. This experiment provided evidence that the *kcnb1* mutant was sensitive to high-intensity light.

**Fig. 4:**
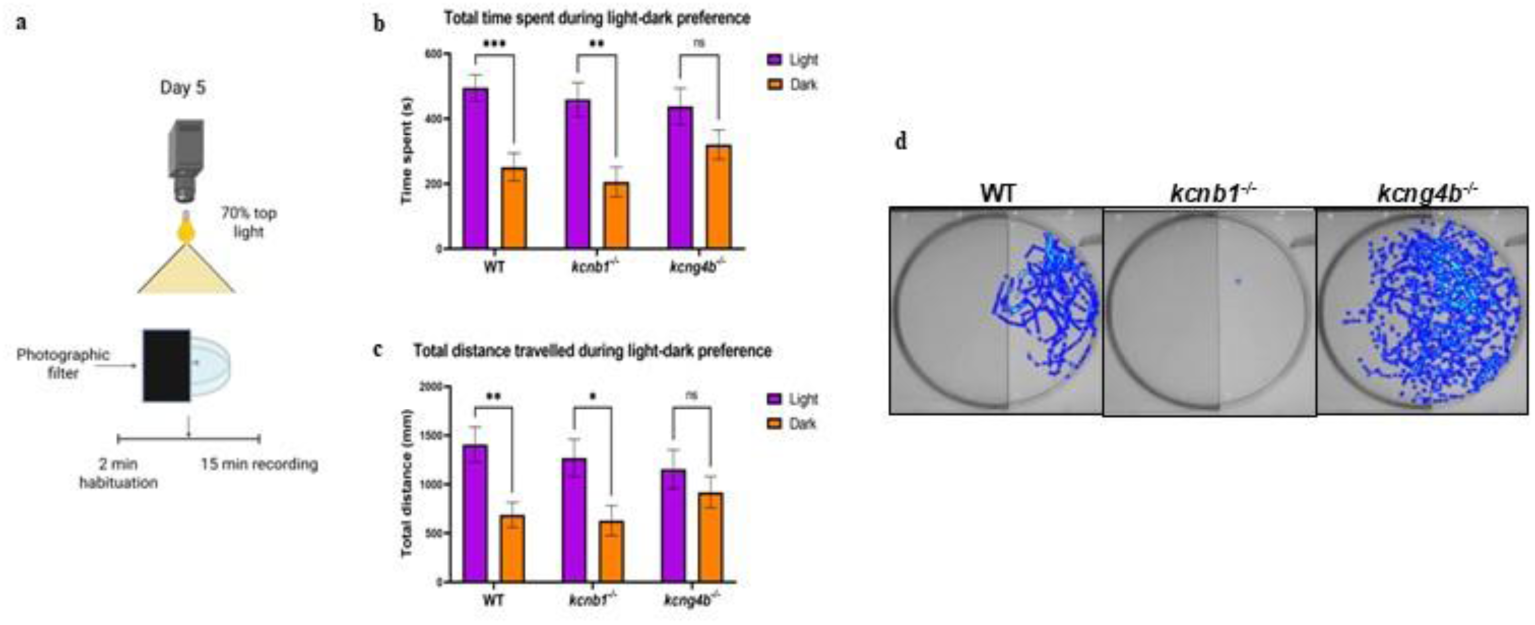
High intensity light induces freezing behavior in the *kcnb1^-/-^* larvae. The light/dark preference test was performed at 5 dpf with 70% top light over a 15-minute period. a) Schematic representation of the light-dark setup. **b) Light zone preference:** The total time spent in light zone is higher in the WT and *kcnb1* mutant larvae, indicating a light preference. In contrast, the kcng4b mutant shows no preference, spending similar time in both zones. **c) Locomotor activity in light zone:** The total distance travelled in the light zone is larger in the WT and *kcnb1* mutant groups. Statistics were calculated using two-way ANOVA followed by Ŝídák’s multiple comparison test. (N=4; n WT=32, n *kcnb1^-/-^*=32, n *kcng4b^-/-^*=30); **d) Freezing behavior:** The heatmaps obtained from Ethovision XT represent the individual trajectory and total distance covered across the arena. Note the freezing behavior of the *kcnb1* mutant larvae upon high-intensity light exposure All data are represented as mean ± SEM. A p-value of ≤0.05 was considered statistically significant.

Based on the results of the light/dark preference test, we performed two additional behavioral tests to check the response of the *kcnb1* mutant to high-intensity light. The first one was an open field test in both dark and high-intensity light (70% top light). The total distance travelled in the dark decreased in the mutants, while the mean velocity of both groups, wild-type and *kcnb1^-/-^*, was similar (Fig. 5A-a and b). However, under high-intensity light, the total distance travelled, and the mean velocity were significantly reduced in the *kcnb1* mutants (Fig. 5A-c and d). Moreover, normal behavior was restored in mutants as soon as the lights turned off at the end of the experiment.

**Fig. 5:**
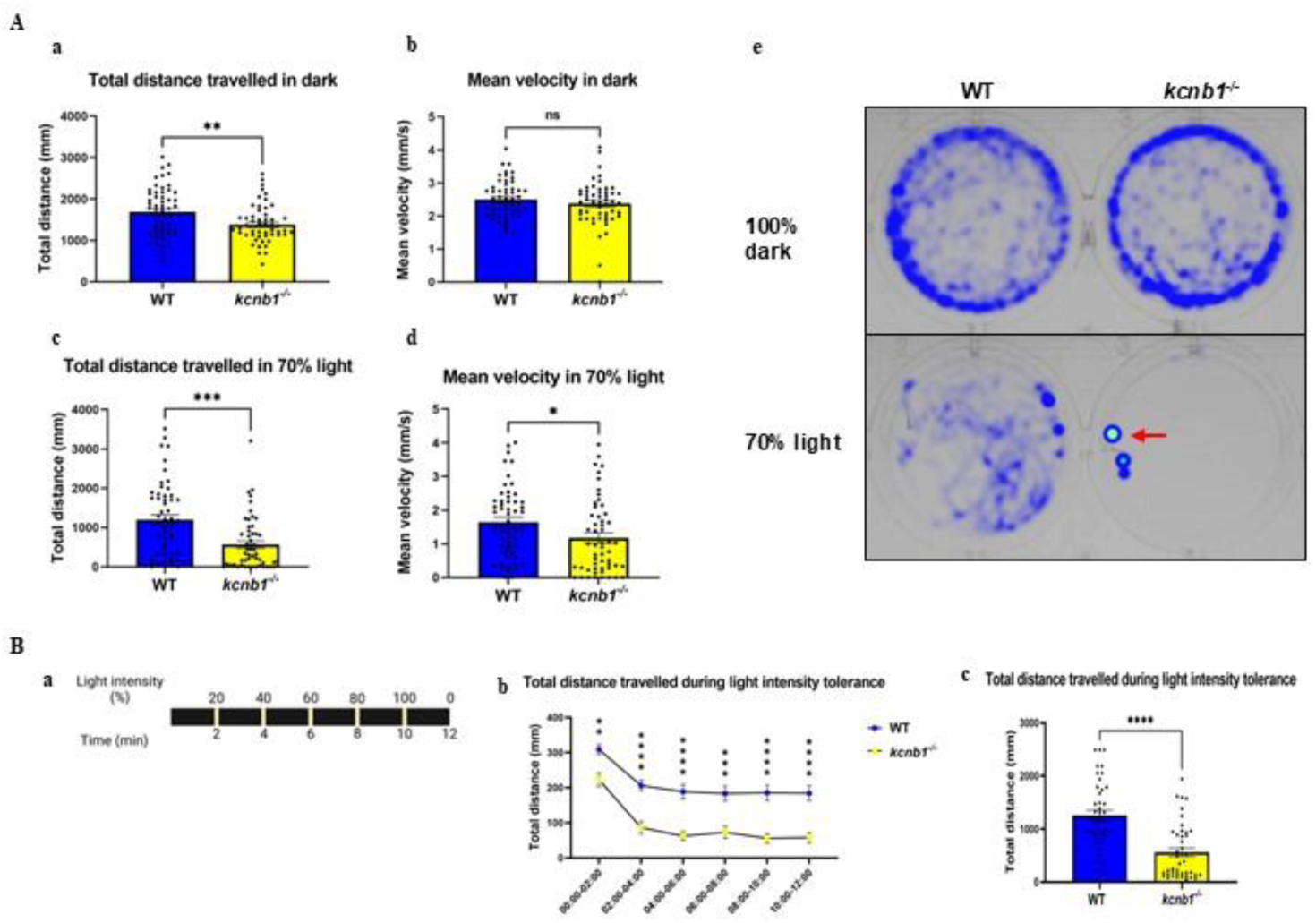
The kcnb1^-/-^ larvae displayed increased photosensitivity to high-intensity light. **A. Open field test under different conditions: a-b) Locomotor activity in dark:** Under dark conditions, the total distance travelled (a) was reduced in the kcnb1 mutant compared to WT. However, mean velocity (b) showed no changes. Statistics for (a) were calculated using unpaired Student’s t-test with Welch’s correction while (b) used a non-parametric Mann-Whitney test. (N=3; n=54); **c-d) Locomotor activity in light:** The locomotor activity of the kcnb1^-/-^ larvae was significantly decreased (c-d) under high-intensity light (70% top light), suggesting sensitivity. Statistics were calculated using a non-parametric Mann-Whitney test. (N=3; n=54); **e) Freezing behavior:** Heatmaps illustrate the total distance travelled by the larva under light and dark conditions. Note the freezing behavior of the *kcnb1* mutant larvae upon high-intensity light exposure with the red arrow indicating the freezing location. **B. Light intensity tolerance test:** a) Schematic representation of the experimental protocol involving a gradual increase in light intensity. b) The total distance covered by the kcnb1 mutant larvae gradually decreased as the light intensity increased. Statistics were calculated using a mixed effect model followed by Ŝídák’s multiple comparison test. (N=3; n=42); c) The overall distance traveled by the kcnb1 mutant larvae is significantly decreased. Statistics were calculated using a non-parametric Mann-Whitney test. (N=3; n=42). All data are represented as mean ± SEM with each dot representing individual larvae. A p-value of ≤0.05 was considered statistically significant.

The second behavioral test checked the photosensitivity threshold by gradually increasing the top light intensity from 0 to 100% every two minutes (Fig. 5-Ba). Both groups showed a decrease in overall locomotor activity at 20% intensity, with a steady decline as the intensity increased, but the decrease was more robust in the mutant (Fig. 5B-b). However, the total distance travelled by the *kcnb1* mutant larvae significantly decreased (Fig. 5B-c). Together, these results show an increased photosensitivity in the *kcnb1* mutant, evidenced by light-triggered decreased mobility and freezing.

### Mutations in the Kv2.1 and Kv6.4 subunits disrupt inhibitory neurotransmission

Quantitative PCR (qRT-PCR) was performed to analyze the transcriptional levels of several genes implicated in epilepsy. The results showed an increase in the expression levels of *fosab* (*c-fos*) in both the mutants, indicating increased brain activity or hyperexcitability (Fig. 6a). Moreover, in the mutants, there was a significant increase in the levels of glutamate decarboxylase 2 (*gad2*), an enzyme responsible for synthesizing the inhibitory neurotransmitter GABA (Fig. 6b). However, there was a significant reduction in the transcript of the key subunit of the GABA-A receptor-gamma-aminobutyric acid type A receptor subunit alpha-1 (*gabra1*) (Fig. 6c). No changes in the transcript level of *bdnf* (brain-derived neurotrophic factor) was observed (Supplementary table 2). These results suggest a disruption in the gene expression related to the inhibitory network, characterized by an upregulation in *gad2* and a downregulation in *gabra1*, alongside elevated *c-fos* levels.

**Fig. 6:**
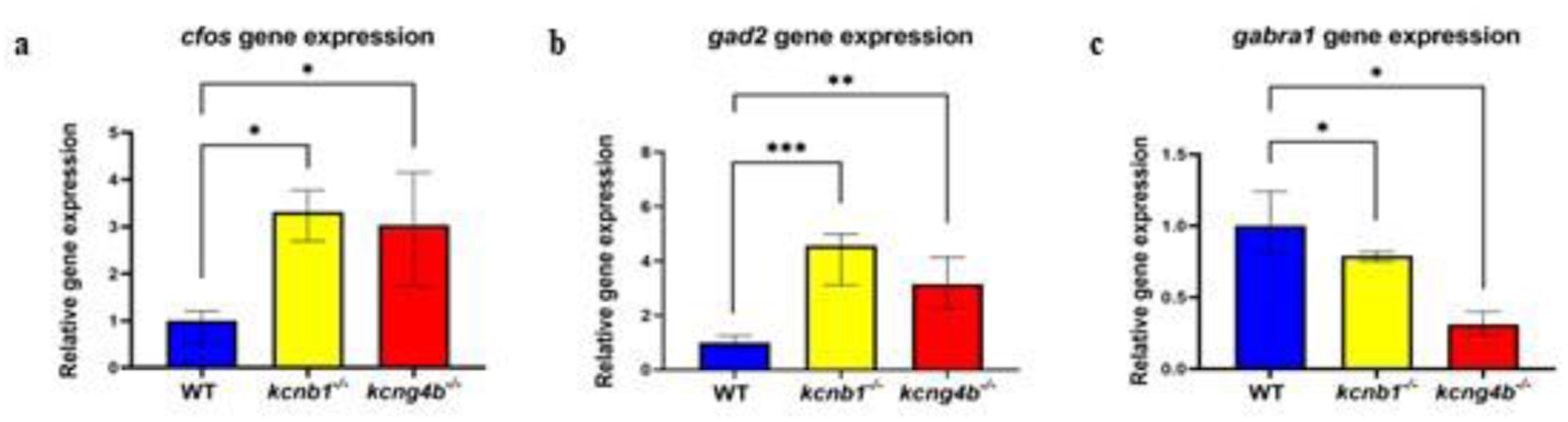
Alteration in transcript level of genes involved in inhibitory neurotransmission. qRT-PCR was performed to analyse the transcript levels of genes involved in epilepsy and inhibitory neurotransmission. **a) Neuronal activity marker:** Increased expression of *c-fos* (immediate early gene) transcript in both *kcnb1^-/-^* and *kcng4b^-/-^*. **b-c) Inhibitory neurotransmission markers:** The gene expression level of *gad2* (glutamate decarboxylase 2), GABA synthesis enzyme (b) is significantly increased in the mutants. The transcript level of *gabra1* (GABA A receptor, subunit alpha-1) was significantly downregulated in both the mutants. Statistics were calculated using one-way ANOVA. All data are represented as mean ± SD with the p-value adjusted to ≤0.05.

Furthermore, transcript level of genes involved in visual perception, *cry2* (cryptochrome circadian regulator 2), and *gnat2* (guanine nucleotide binding protein (G protein), alpha transducing activity polypeptide 2), were upregulated in the *kcnb1* but not in the *kcng4b*. C*ry1a* (cryptochrome circadian regulator 1a), a gene associated with the circadian clock, was significantly downregulated in the *kcnb1*, whereas its expression was significantly upregulated in the *kcng4b* (Supplementary table 2). The changes in the transcript level involved in visual perception suggest that Kv2.1 subunits may play a role in vision.

### Kv2.1 deficiency does not affect the morphology of the retina or the retinotectal projections

Since both qRT-PCR and behavioral tests showed changes in the light-evoked response of the *kcnb1* larvae, we next performed antibody staining to assess the retinal morphology. Nuclear staining did not show any morphological defects in the retinal cell layers in the *kcnb1* mutant (Supplementary Fig. 2A-a and d). The labelling of cone photoreceptors and Müller glia appeared normal in both genotypes (Supplementary Fig. 2A-b to f).

Because there were no visible morphological defects in the retinal layers, we performed DiI staining to trace the retinotectal projections at 5 dpf. The anterograde tracing of retinotectal projections showed normal projections in the *kcnb1* mutant (Supplementary Fig. 2B-g and h). These results together show that neither the morphology of the retina, nor the integrity of retinotectal projections were compromised in the *kcnb1* mutants.

## DISCUSSIONS

In this study, we characterized the effects of LOF of *kcnb1* and *kcng4b* on zebrafish behavior and circuit function. Our primary finding is a convergence of neural phenotypes: both the mutants exhibited increased susceptibility to PTZ-induced seizures, upregulation of the immediate early gene *c-fos*, and altered inhibitory neurotransmission. Unexpectedly, we also identified an interesting phenomenon, i.e., *kcnb1* loss led to an abnormal light-evoked behavioral response, manifested by reduced locomotor activity and freezing under high-intensity light, despite intact retinal and retinotectal morphology. Together, these findings reveal shared roles of *kcnb1* and *kcng4b* in the inhibitory network and a unique role for *kcnb1* in sensory processing.

Previous study on the zebrafish *kcnb1* LOF mutant has shown hyperexcitability and GABA dysregulation, characterizing features of KCNB1-related encephalopathies in humans (44). Our study extends these findings by analyzing both α subunit (*kcnb1*) and modulatory subunit (*kcng4b*) mutants in parallel and by identifying a sensory component associated with *kcnb1* loss. Even though the developmental studies demonstrated the antagonistic effect of the Kv2.1 subunits during development of hollow organs (29–32), our data show that their neural functions converge at the level of circuit excitability and stability. These results highlight a context-dependent link between Kv2.1-Kv6.4 subunits across development and neural processes.

Even though initially the mutants are hypoactive, the upregulated *c-fos* levels, spontaneous brain excitability, and susceptibility to lower concentrations of PTZ-induced seizures collectively reveal an underlying state of neuronal hyperexcitability. This evident physiological hyperexcitability, but lack of spontaneous behavioral manifestations, suggests that these mutants can still serve as a valuable model for screening compounds that correct the fundamental hyperexcitability, even if it does not fully mimic the human DEE seizure phenotype.

Furthermore, the data show a disrupted inhibitory system shared by both mutants. Kv2.1 channels are known to regulate neuronal excitability not only through delayed-rectifier potassium currents but also by clustering and interaction with other ion channel subunits in an activity-dependent manner (45,46). Therefore, losing either *kcnb1* or *kcng4b* likely alters the way these ion channel complexes work, leading to weakened inhibitory control across the neural circuit. This state of central hyperexcitability is further supported by the contradictory changes observed in the transcription levels of inhibitory neurotransmission markers. We found a significant increase in GABA-synthesizing enzyme *gad2*, which was also observed by other authors (44), which we interpret as a compensatory mechanism, where the central nervous system attempts to suppress excitability by increasing the supply of the inhibitory neurotransmitter GABA. However, these efforts appear ineffective due to a simultaneous downregulation of the GABA-A receptor subunit *gabra1*. The depletion of postsynaptic GABA-A receptors represents a critical constraint, suggesting that despite an elevated level of GABA, the postsynaptic inhibitory infrastructure is severely compromised. This leads to a net reduction in inhibitory neurotransmission, thereby maintaining the hyperexcitable state confirmed by the *c-fos* and electrophysiology results. This interpretation aligns with the findings in the *gabra1^-/-^* zebrafish model, where loss of this subunit leads to a drastic reduction in inhibitory synapses and network complexity (47).

In addition to the observed hyperexcitability, *kcnb1* loss uniquely leads to decreased locomotor responses and freezing under bright illumination, suggesting an abnormal photic response rather than photosensitive seizures. In absence of retinal or retinotectal defects the impairment may arise not from neuroanatomical deficiency, but rather from altered signaling properties within the visual pathway. The severe behavioral freezing suggests a fundamental failure in the transduction or transmission of the light-evoked signal at the photoreceptor level. Although *kcng4b* has shown overlapping expression with *kcnb1* in the eye (29), its loss does not recapitulate the acute light-evoked deficit. Prior mouse models have elegantly shown that Kv2.1 and Kv8.2 loss disrupts the retinal dark current and phototransduction kinetics (33,34,48), yet our findings provide the first evidence linking Kcnb1 to a specific, quantifiable, light-evoked behavior in a whole, developing vertebrate organism. This acute behavioral signature effectively validates the known functional disruption of the Kv2.1/Kv8.2 channel in a whole-animal context.

The functional nature of this light-evoked response is further highlighted by the observation that the *kcnb1* larvae locomotor activity normalizes as soon as the light is switched off. This rapid recovery confirms that the defect is a consequence of acute, dysregulated electrical signaling in the visual pathway, rather than a long-lasting or permanent structural or developmental defect. This faulty signal originating from the retina could be amplified by the pre-existing state of hyperexcitability in the central nervous system, confirmed by elevated *c-fos* transcript levels and the LFP recordings showing increased spontaneous activity. Crucially, this central hyperexcitability acts as an amplifier, converting the subtle retinal signaling error into the severe, dramatic behavioral freezing we observe under high-intensity light, which represents the basis for the ‘two-hit’ model. Furthermore, whereas WT and *kcnb1* larvae display a natural preference for the light zone, the *kcng4b* mutant exhibits a complete lack of preference (Fig. 4b-c), suggesting a distinct, non-acute disruption of the light/dark processing pathway.

Finally, the transcriptional changes observed in genes related to light signaling, specifically the upregulation of *cry2* and *gnat2* and the downregulation of *cry1a* in the *kcnb1* mutant, provide molecular evidence for *kcnb1’s* role in the visual circuit. Notably, the *kcng4b* mutant showed significant upregulation of the *cry1a* transcript level, highlighting a transcriptional divergence in the sensory/circadian network. While these changes may reflect long-term compensatory responses to altered light input or an impact on the circadian network, their specific dysregulation in the *kcnb1* mutant (but not in *kcng4b*) reinforces the hypothesis that *kcnb1* plays a unique, *kcng4b*-independent role in modulating the photic response pathway. Together, these findings suggest that Kv2.1 contributes to the stability of visual circuits by regulating both essential photoreceptor signaling and central inhibitory tone.

Overall, our findings highlight the essential contribution of Kv2.1 subunits to inhibitory homeostasis and sensory processing. The parallel phenotypes of *kcnb1* and *kcng4b* mutants reveal shared pathways that may underlie the variability observed in human *KCNB1*-related disorders. The zebrafish model provides a flexible system to dissect these mechanisms further, including subunit-specific interactions, developmental timing of the Kv2.1 function, and circuit-level consequences of the channel imbalance. Future studies combining genetic, electrophysiological, and pharmacological approaches will be crucial to clarify how Kv2.1 channel diversity modulates excitability across neural systems.

## Funding declaration

Korzh V acknowledges support from the Opus grant of the National Science Centre (NCN), Poland (2020/39/B/NZ3/02729).

**Consent to publish:** not applicable.

**Consent to participate:** not applicable.

## Ethics declaration

Zebrafish (*Danio rerio*) were maintained according to established protocols (Westerfield, 2007) in the Zebrafish Core Facility at the International Institute of Molecular and Cell Biology in Warsaw (licensed breeding and research facility, PL14656251, registry of the District Veterinary Inspectorate in Warsaw; 064 and 051: registry of the Ministry of Science and Higher Education). All the experiments with zebrafish embryos, larvae and adults were performed in accordance with the European Communities Council Directive (63/2010/EEC).

## Data availability declaration

the materials and reagents described in the paper are available from authors upon request.

## Author contribution declaration

Ruchi P. Jain: Conceptualization, Methodology, Investigation, Writing – original draft, Project administration, Data curation. R. Rosa Amini: Methodology, Investigation. Lukasz Majewski: Methodology, Investigation. Vladimir Korzh: Writing – review & editing, Validation, Supervision, Project administration, Funding acquisition.

## Declaration of generative AI and AI-assisted technologies in the writing process

During the preparation of this work the author(s) used Grammarly and Quillbot to improve language and readability. After using this tool/ service, the authors reviewed and edited the content as needed and took full responsibility for the content of the publication.

## Acknowledgements

The authors are thankful to Prof. Jacek Kuznicki and all members of the Laboratory of Neurodegeneration (IIMCB in Warsaw) for fruitful communication, the Microscopy and Zebrafish Core Facilities (IIMCB in Warsaw) for expert technical help and fish maintenance.

---

**Supplementary Fig. 1: High intensity light induces freezing behavior in the kcnb1^-/-^ larvae.**

The light/dark preference test was performed at 5 dpf with 70% top light over a 15-minute period. a) Mean velocity between the WT and mutants (*kcnb1^-/-^* and *kcng4b^-/-^*) larvae remains unaltered; b) Zone transition frequency for WT and *kcnb1^-/-^* is higher in light than dark, but *kcng4b^-/-^* shows no changes. Statistics were calculated using two-way ANOVA followed by Ŝídák’s multiple comparison test. (N=4; n WT=32, n *kcnb1^-/-^* =32, n *kcng4b^-/-^*=30).

All data are represented as mean ± SEM. A p-value of ≤0.05 was considered statistically significant.

**Supplementary Fig 2: No morphological defects in the retina or retinotectal projections.**

**A. Retinal cross-section analysis:** Transverse retinal section from 5 dpf WT (a-c) and *kcnb1^-/-^* (d-f) larvae show no morphological defects. Nuclear staining (Hoechst, blue) highlights the retinal cell layers (a,d). Cone photoreceptors (red) are labelled by anti-gnat2 (b,e), and Müller glia (green) are labelled by anti-glutamine synthetase (c.f). All markers and layers appear normal between the genotypes. MG-Müller glia.

**B. Retinotectal projection Integrity:** Anterograde labelling of the retina was performed using DiI (1,1’-Dioctadecyl-3,3,3’,3’-Tetramethylindocarbocyanine Perchlorate) in 5 dpf WT (g) and *kcnb1^-/-^* (h) larvae. The retinotectal projections in the optic tectum appear normal, with no defects in *kcnb1* mutant compared to the wildtype. OT-optic tectum, E-eye.

