## Supplementary figures and images for "Kv2.1-Kv6.4 subunits deficiency impairs inhibitory signaling and visual circuit dynamics in zebrafish"

### Supplementary Figures 1-2

**a**

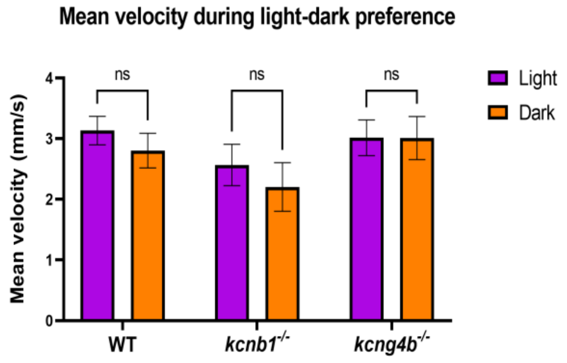


**b**

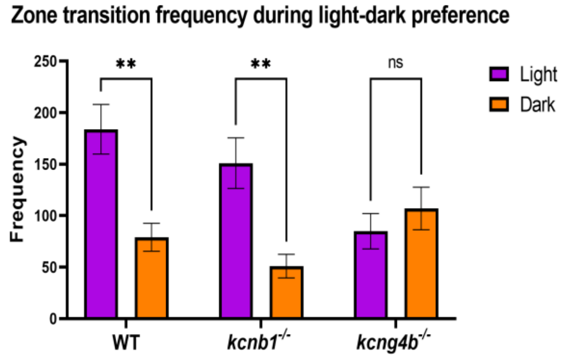


**Supplementary Fig. 1**

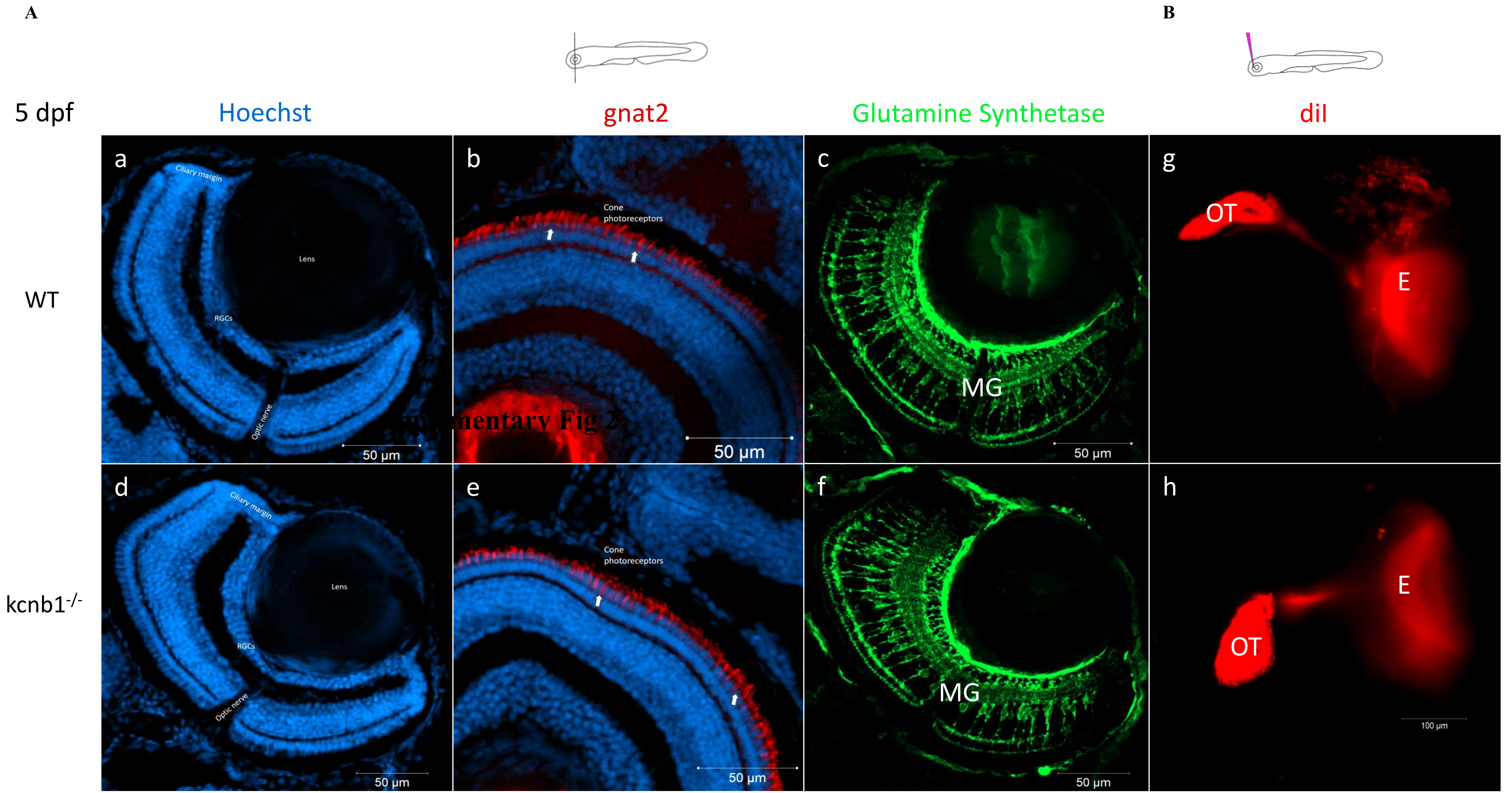


**Supplementary Fig 2**
