## Supplementary Table 1 for "Kv2.1-Kv6.4 subunits deficiency impairs inhibitory signaling and visual circuit dynamics in zebrafish"

| **Genes** | **qRT-PCR primers, 5'-> 3'** |
| --- | --- |
| *bdnf* | TGACAGTATTAGCCAGTG  CATTGCGAGTTATAGTGC |
| *fosab/c-fos* | GATGGCTGCTGCGAAATGC  CGGCGAGGATGAACTCTAACC |
| *gad2* | CATTGAAGCCAAGCAGAAG  GAACCAGAAGAGCAGAGC |
| *gabra1* | TGTGTGGAGTTTGTGTGAAAGC  TCATTTGCGTAGCGAGGAGG |
| *cry1a* | TTCTTCCAGCAGTTCTTCC  ATCGCCGTATGTAGTCTCC |
| *cry2* | CTGTTGGATGCGGACTGG  CTCGTAGATGTAGCGGTTAGG |
| *gnat2* | CCACTTGTCCACCTCCAC  CTTCCTTATCGGCATCTTCC |
