## Supplementary Table 2 for "Kv2.1-Kv6.4 subunits deficiency impairs inhibitory signaling and visual circuit dynamics in zebrafish"

| **Genes** | **Expression level ratio,**  ***kcnb1* / wt, SD**  **5dpf** | **p-value** | **Expression level ratio,**  ***kcng4b* / wt, SD**  **5dpf** | **p-value** |
| --- | --- | --- | --- | --- |
| *bdnf* | 0.97±0.15 | ns | 0.74±0.16 | ns |
| *fosab* | 3.31±0.15 | 0.05 | 3.03±0.26 | 0.05 |
| *gad2* | 4.56±0.25 | 0.001 | 3.15±0.1 | 0.01 |
| *gabra1* | 0.79±0.05 | 0.05 | 0.31±0.08 | 0.05 |
| *cry1a* | 0.23±0.02 | 0.006 | 6.35±0.28 | 0.001 |
| *cry2* | 1.84±0.43 | 0.05 | 0.83±0.14 | ns |
| *gnat2* | 1.61±0.27 | ns | 0.46±0.11 | ns |
